# Evaluating longitudinal ecological models linking scientific production to population-level indicators: a global case study in mental health research

**DOI:** 10.64898/2026.05.09.723946

**Authors:** Andy A. Acosta-Monterrosa, David A. Hernández-Paez, Fabriccio J. Visconti-Lopez, Sulaiman Kalokoh, Ivan David Lozada-Martinez

## Abstract

**Background:** Quantifying the alignment between scientific production and population-level indicators remains a persistent methodological challenge in health research evaluation. While longitudinal ecological models have been increasingly used to explore associations between research output and societal outcomes, their feasibility, interpretability, and structural limitations have not been systematically examined.

**Methods:** We conducted a longitudinal ecological meta-research analysis integrating global bibliometric data on mental health publications with country-level indicators of mental disorders, mental health infrastructure, and subjective well-being. Analyses were stratified by World Bank income groups and implemented using a three-step framework comprising income specific linear regression models, random-effects meta-analyses, and meta-regressions to assess association patterns, heterogeneity, and potential moderators.

**Results:** Scientific production was highly concentrated in high-income countries. Income-stratified regression models revealed divergent association patterns across contexts, with inverse associations observed in higher income groups and predominantly positive coefficients in low-income countries. Meta-analyses showed extreme between-group heterogeneity for most indicators, yielding largely attenuated pooled estimates. Only one subjective well-being indicator retained a significant pooled association.

**Conclusions:** Longitudinal ecological models linking scientific production to population-level indicators can identify broad association patterns and structural asymmetries but are strongly constrained by contextual heterogeneity and data availability.

## 1. Introduction

Over the last decades, scientific production in health-related fields has expanded at an unprecedented pace. Advances in bibliometrics and research analytics have made it increasingly feasible to quantify this growth, track its geographic distribution, and characterize its thematic evolution [1]. However, despite the availability of increasingly granular publication data, a fundamental methodological question remains unresolved: to what extent can scientific production be meaningfully linked to population-level indicators of health, well-being, and societal development over time [2,3].

Attempts to address this question have traditionally relied on indirect proxies of research performance, such as citation-based metrics, journal impact indicators, or funding volume [4]. While useful for characterizing academic influence, these measures offer limited insight into whether scientific activity aligns with, precedes, or plausibly contributes to changes in real-world outcomes [4]. Bridging this gap requires methodological approaches capable of integrating heterogeneous data streams (bibliometric, epidemiological, and socio-developmental) across long temporal horizons and diverse geopolitical contexts [5]. Yet, the feasibility, interpretability, and limitations of such approaches remain insufficiently examined.

Ecological and longitudinal study designs have emerged as one possible strategy to explore these relationships at scale [6,7]. By aggregating publication outputs and population-level indicators over time, these models allow for the exploration of temporal associations while accommodating global heterogeneity [6,7]. Variants of this approach have been applied in multiple domains, including health services research, patient safety, quality improvement, and meta-research [6-8]. Nonetheless, these applications often remain domain-specific, and few studies have explicitly interrogated the methodological performance of the approach itself, particularly its sensitivity to data sparsity, structural heterogeneity between countries, and the uneven availability of surveillance indicators.

Mental health research represents a particularly instructive empirical context in which to examine these methodological challenges [9]. On the one hand, mental health has experienced substantial growth in scientific output, especially in high-income settings, generating a large and diverse bibliometric corpus [9]. On the other hand, population-level mental health indicators, ranging from disorder prevalence and disability-adjusted life years to infrastructure and subjective well-being measures, are characterized by fragmented coverage, inconsistent reporting, and substantial missingness across countries and income groups [10]. This combination of abundant research output and uneven outcome data creates a natural stress test for longitudinal ecological models that aim to link science to population-level metrics.

Importantly, the objective of such analyses is not to infer causal effects of research on health outcomes, nor to evaluate the effectiveness of mental health interventions at the population level. Rather, the methodological question is whether longitudinal ecological frameworks can generate stable, interpretable, and context-sensitive associations between scientific production and population-level indicators, and under what conditions these associations break down. Addressing this question requires explicit attention to model assumptions, data structure, heterogeneity, and the limits of inference inherent to aggregate analyses.

In this study, we evaluate the methodological feasibility, strengths, and limitations of linking global scientific production to population-level indicators using a longitudinal ecological design. Using mental health research as a case study, we integrate large-scale bibliometric data with country-level indicators of mental disorders, mental health infrastructure, and subjective well-being across income groups. A three-step analytical framework (comprising linear regression, random-effects meta-analysis, and meta-regression) was employed to assess the consistency, heterogeneity, and potential moderators of observed associations over time.

By focusing on the methodological behaviour of these models rather than on substantive outcome interpretation, this work aims to contribute empirical evidence to ongoing discussions in meta-research and research evaluation [11]. Specifically, it seeks to clarify what longitudinal ecological approaches can and cannot reveal about the alignment between scientific activity and population-level indicators, and to identify structural constraints, such as data availability and income-related disparities, that shape their interpretability. These insights are intended to inform future methodological developments in the assessment of research coherence, impact, and accountability at the global level.

## 2. Methods

### 2.1. Study design

This study was conceived as a longitudinal ecological meta-research analysis designed to examine the methodological behaviour of models that link scientific production to population-level indicators over time. Rather than aiming to estimate causal effects or intervention effectiveness, the analytical framework was structured to evaluate whether such models can generate stable, interpretable, and context-sensitive associations when applied at a global scale [12].

### 2.2. Data sources and selection

#### 2.2.1. Scientific production data

Global scientific production related to mental health was identified through a systematic search strategy detailed in the **Supplementary Material 1**. The search retrieved 438,837 records. After restricting the dataset to document types with complete bibliometric metadata, 386,671 publications were retained for analysis. Inclusion required the availability of publication year, first author’s country affiliation, journal impact indicators (H-index and quartile), total citation counts, and access type.

Country affiliations were classified according to the World Bank income group categories (low-, lower-middle-, upper-middle-, and high-income) [13]. This stratification was applied consistently across all analytical stages to preserve comparability between bibliometric and population-level indicators. The use of publication volume as the primary exposure variable reflects a pragmatic methodological choice: it represents a reproducible, transparent, and longitudinally stable proxy of scientific activity, while acknowledging that it captures quantity rather than content, quality, or translational uptake.

#### 2.2.2. Population-level indicators

Country-level indicators were obtained from publicly accessible repositories, including the World Health Organization Global Health Observatory [14], the World Bank [15], Our World in Data [16], and the Institute for Health Metrics and Evaluation [17]. Data extraction was conducted through public application programming interfaces when available; otherwise, manual retrieval procedures were applied.

A total of 24 indicators related to mental health and subjective well-being were included. For analytical clarity, indicators were grouped into three domains: 1) mental disorders; 2) mental health infrastructures; and 3) subjective well-being. A full description of indicators, definitions, and aggregation methods is provided in **Supplementary Table 1**.

To enable cross-country comparisons while accounting for population size, indicators were aggregated using population-weighted averages at the income-group level. This aggregation strategy reflects an explicit methodological trade-off: it reduces within-group variability while increasing robustness against country-level data sparsity, a recurrent limitation in global mental health surveillance.

### 2.3. Statistical Analysis

A three-step analytical framework was adopted, comprising linear regressions, meta-analyses, and meta-regressions.

#### 2.3.1. First step

Linear regression models were used to examine associations between publication volume and population-level indicators. Analyses were stratified by income group to account for structural differences in research capacity, health systems, and data availability.

Depending on the indicator, publication volume was specified as either the independent or dependent variable, reflecting the bidirectional nature of ecological associations and avoiding implicit assumptions regarding directionality. For indicators related to mental disorders, regression coefficients (β_1_) were normalized using z-score standardization to facilitate comparison across measures with different scales. For infrastructure and subjective well-being indicators, raw β_1_ coefficients were retained to preserve interpretability [18].

Given that this analysis is part of a broader research program, only results pertaining to mental health indicators are presented in the main manuscript. Full regression outputs for all indicators are provided in **Supplementary Material 2**.

#### 2.3.2. Second step

To synthesize association estimates across income groups, random-effects meta-analyses were conducted for each indicator. This approach was selected to explicitly model between-group heterogeneity rather than assume a common underlying effect.

Pooled estimates were calculated by integrating income-group-specific regression coefficients and their standard errors. Between-group variance components were estimated using restricted maximum likelihood methods. Heterogeneity was assessed using the I^2^ statistic and Cochran’s Q test [18]. Complete meta-analytic results are reported in **Supplementary Material 3**. The use of meta-analysis in this context is methodological rather than inferential. Pooled estimates are interpreted as summaries of association patterns across structurally distinct contexts, not as global effect sizes.

#### 2.3.3. Third step

Meta-regression analyses were performed to explore potential moderators of between-group heterogeneity identified in the meta-analyses. Candidate moderators included all available population-level indicators as well as selected bibliometric characteristics.

To avoid overfitting and preserve interpretability, only moderators explaining the largest proportion of heterogeneity for each mental health indicator are reported in the main text [18]. Full meta-regression outputs are provided in **Supplementary Material 4**.

Importantly, the meta-regression stage was not intended to exhaustively explain heterogeneity, but to assess whether systematic patterns could be identified beyond random variation, given the structural constraints of the available data [18].

### 2.4. Missing data and structural limitations

The availability of population-level mental health indicators varied substantially across income groups and over time. Missingness was quantified descriptively for each indicator and income group, and no imputation procedures were applied. This decision reflects a methodological stance: missing data patterns were treated as informative features of the global mental health surveillance landscape rather than as technical nuisances to be corrected [19].

Indicators with insufficient data coverage within specific income groups were excluded from regression analyses for those strata, and such exclusions are reported transparently in the results.

### 2.5. Statistical software and reproducibility

All analyses were conducted using R (version 4.5.0). Annotated scripts and documentation supporting the analytical workflow are publicly available at https://doi.org/10.5281/zenodo.18417489, ensuring full reproducibility and facilitating methodological scrutiny.

## 3. Results

### 3.1. Descriptive bibliometric patterns

Global scientific production related to mental health, when stratified by World Bank income groups, was markedly concentrated in high-income countries. Publications originating from this group accounted for 81.2% of the total output identified by the search strategy **(Table 1)**, whereas low-income countries contributed only 0.15% of the overall production. The earliest mental-health-related publication included in the dataset dates back to 1886; however, representation from all income groups was not observed until 1968 **(Figure 1)**. From that point onward, publication volume increased steadily across all strata.

**Table 1.**
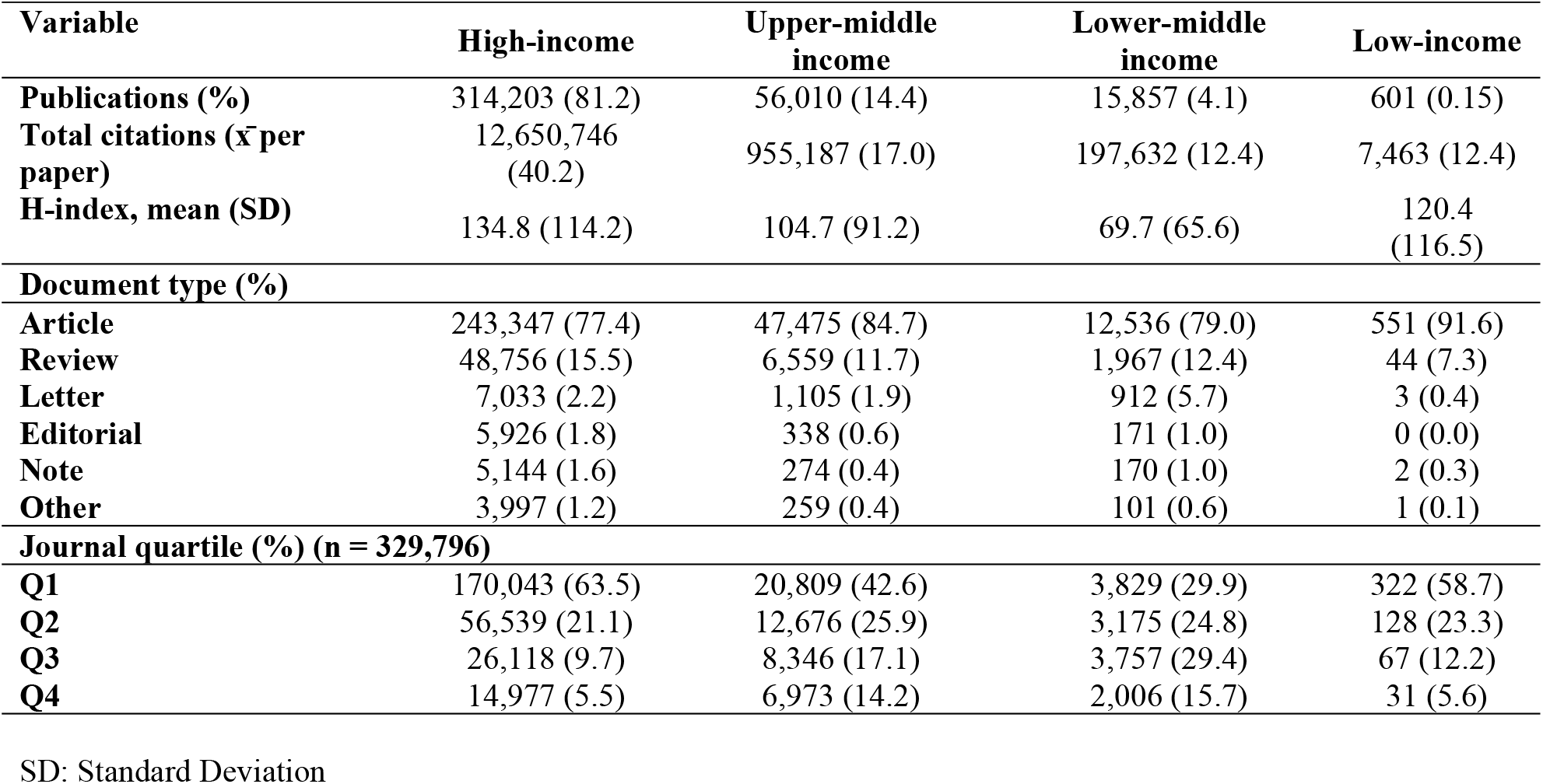
General characteristics of articles on mental health by region (N = 386,671)

**Figure 1.**
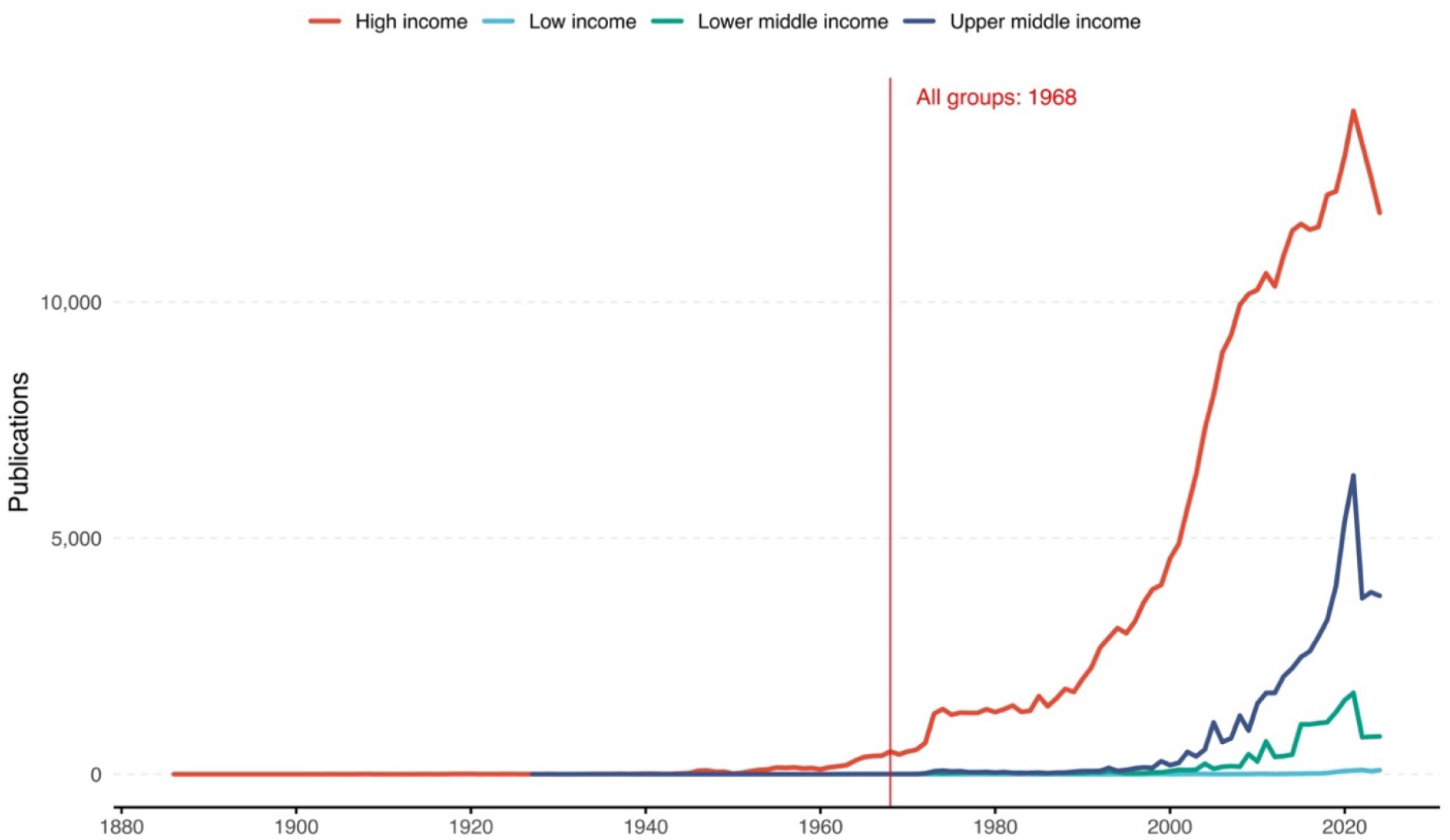
Annual trends in mental health-related publications by income (1886-2024)

Beyond volume, substantial disparities were also observed in citation patterns. Articles from high-income countries accumulated a mean of 40.2 citations per paper **(Table 1)**, considerably exceeding the averages observed in upper-middle-, lower-middle-, and low-income groups. Original research articles constituted the predominant document type across all income categories, although their relative proportion was highest in low-income countries (91.6%). Review articles represented the second most common format in all strata.

Journal impact profiles further reflected income-related asymmetries. High-income countries published the majority of their articles in first-quartile journals (63.5%), followed by second-quartile outlets (21.1%). A similar pattern was observed in upper-middle-income countries, whereas lower-middle-income countries showed a more even distribution across quartiles. Despite their limited output, publications from low-income countries were also predominantly placed in first- and second-quartile journals.

### 3.2. Income-stratified regression and meta-analytic findings

Within the longitudinal ecological framework, linear regression models were used to characterize associations between publication volume and population-level mental health indicators across income groups. For indicators related to mental disorders, including anxiety, bipolar disorder, eating disorders, and schizophrenia, measured as prevalence or disability-adjusted life years (DALYs), inverse associations were consistently observed in high-, upper-middle-, and lower-middle-income countries, with the strongest coefficients generally detected in high-income settings **(Figure 2a)**.

**Figure 2.**
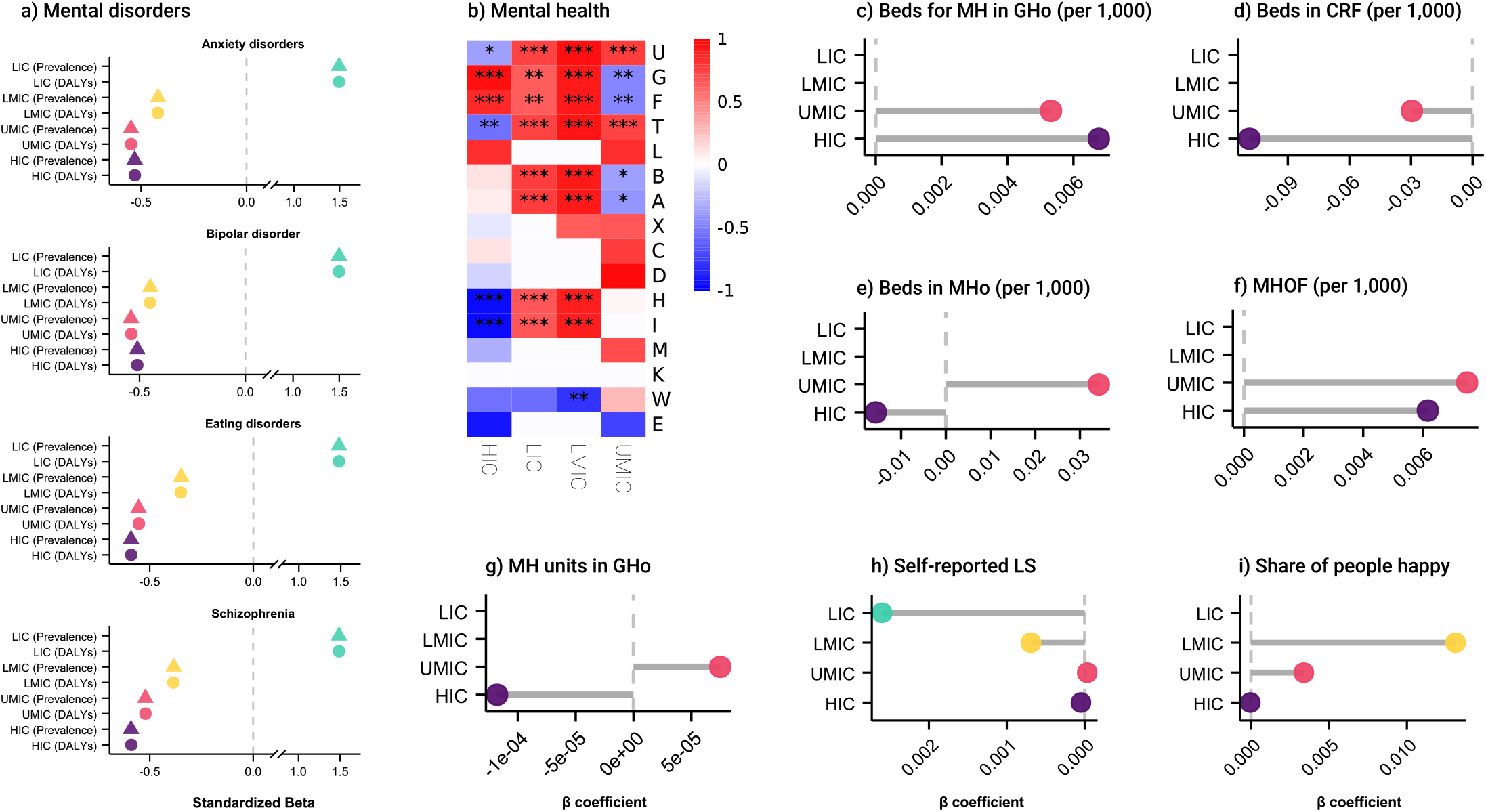
Associations between mental-health-related publication volume and indicators of mental health across income groups. a) Display z-score normalized β_1_ coefficients for grouped indicators related to mental disorders. b) Heatmap showing z-score normalized β_1_ coefficients for the association between the volume of mental-health-related publications and fifteen indicators on mental health across high-income countries (HICs), upper-middle-income countries (UMICs), lower-middle-income countries (LMICs), and low-income countries (LICs). Asterisks within cells indicate statistical significance (*p < 0.05, **p < 0.01, ***p < 0.001), while asterisks next to variable abbreviations denote indicators included as independent variables in the regression models. Full variable names and corresponding abbreviations are provided in Supplementary Table 1-4. c-i) Regression coefficients (β_1_) representing associations between publication volume and the number of beds for mental health (MH) in general hospitals (GHo) c), number of beds in community residential facilities (CRF) d), number of beds in mental hospitals (Mho) e), mental health outpatient facilities (MHOF) f), MH units in GHo g), self-reported life satisfaction (LS) h) and share of people who say they are happy i) across income groups.

In contrast, regression coefficients estimated for low-income countries were predominantly positive for the same indicators, indicating that higher publication volumes were associated with higher estimated prevalence and DALYs within this stratum **(Figure 2a-b)**. These opposing directions across income groups underscore substantial contextual heterogeneity in the associations generated by the models.

For indicators related to mental health infrastructure, regression analyses could not be performed in low- and lower-middle-income countries due to insufficient data availability. Among the ten infrastructure-related models estimated in the remaining income groups, none yielded statistically significant associations between publication volume and infrastructure indicators **(Figure 2c-g)**. A similar pattern was observed for subjective well-being indicators **(Figure 2h-i)**, with the exception of self-reported life satisfaction in lower-middle-income countries, where a small but statistically significant negative coefficient was identified (β_1_ = -0.00068, p < 0.01).

When association estimates were synthesized using random-effects meta-analysis, pooled effects were largely attenuated. Across all indicators examined, only the proportion of individuals reporting happiness retained a statistically significant pooled coefficient (0.0021; 95% CI: 0.002-0.006), accompanied by moderate heterogeneity (I^2^ = 72). All other indicators exhibited extreme between-group heterogeneity (I^2^ values approaching 100%), resulting in non-significant pooled estimates **(Table 2)**.

**Table 2.** Meta-analysis results of global health indicators used as dependent or independent variables in linear models related to number of publications.

| Indicator | Coefficient | (95% CI) | Variable | I<sup>2</sup> | Q |
| --- | --- | --- | --- | --- | --- |
| **Mental disorders** |  |  |  |  |  |
| Anxiety disorders (DALYs) | 0.25 | (-0.23 - 0.72) | dep | 99.9 | 151.78* |
| Anxiety disorders (Prevalence) | 2.03 | (-1.83 - 5.89) | dep | 99.9 | 148.74* |
| Bipolar disorder (DALYs) | 0.04 | (-0.04 - 0.11) | dep | 99.9 | 140.09* |
| Bipolar disorder (Prevalence) | 0.16 | (-0.18 - 0.21) | dep | 99.9 | 143.68* |
| Eating disorders (DALYs) | 0.02 | (-0.02 - 0.05) | dep | 99.9 | 213.23* |
| Eating disorders (Prevalence) | 0.08 | (-0.07 - 0.24) | dep | 99.9 | 216.39* |
| Schizophrenia (DALYs) | 0.07 | (-0.05- 0.18) | dep | 99.9 | 285.57* |
| Schizophrenia (Prevalence) | 0.10 | (-0.07 - 0.27) | dep | 99.9 | 302.53* |
| **Infrastructure** |  |  |  |  |  |
| Beds in mental hospitals (per 100,000) | -0.07 | (-0.14 - 0.01) | dep | 68.4 | 3.17 |
| **Subjective well-being** |  |  |  |  |  |
| Self-reported life satisfaction | -0.0004 | (-0.0011 - 0.0003) | dep | 99.3 | 20.66* |
| Share of people who say they are happy | 0.0021 | (0.002 - 0.006) | dep | 72.0 | 6.66* |

DALYs: Disability-Adjusted Life Years; CI: Confidence Interval.

Meta-regression analyses did not identify moderators that meaningfully explained the observed heterogeneity for any of the mental health indicators included. Although selected bibliometric and population-level variables were explored, none accounted for a substantial proportion of the between-group variance **(Supplementary Material 4)**.

### 3.3. Coverage and completeness of mental health indicators

Assessment of data availability revealed pronounced variability in the temporal coverage of mental health indicators across income groups. When examined over the period 2000-2024, most indicators were measured sporadically, resulting in high proportions of missing values **(Table 3)**. Infrastructure-related indicators and government expenditure measures were particularly affected, with missingness exceeding 90% in several income strata.

**Table 3.** Percentage of missing values of indicators related to mental health, yearly, from 2000-2024.

| Indicator | High income | Upper middle income | Lower middle income | Low income |
| --- | --- | --- | --- | --- |
| Anxiety disorders (DALYs) | 12.0% | 12.0% | 12.0% | 13.0% |
| Anxiety disorders (Prevalence) | 12.0% | 12.0% | 12.0% | 13.0% |
| Beds for mental health in general hospitals (per 100,000) | 84.0% | 88.0% | 88.0% | 95.7% |
| Beds in community residential facilities (per 100,000) | 84.0% | 88.0% | 88.0% | 95.7% |
| Beds in mental hospitals (per 100,000) | 84.0% | 88.0% | 88.0% | 91.3% |
| Bipolar disorder (DALYs) | 12.0% | 12.0% | 12.0% | 13.0% |
| Bipolar disorder (Prevalence) | 12.0% | 12.0% | 12.0% | 13.0% |
| Eating disorders (DALYs) | 12.0% | 12.0% | 12.0% | 13.0% |
| Eating disorders (Prevalence) | 12.0% | 12.0% | 12.0% | 13.0% |
| Government expenditures on mental health (% of health expenditures) | 96.0% | 96.0% | 96.0% | 95.7% |
| Mental health day treatment facilities (per 100,000) | 92.0% | 88.0% | 92.0% | 100.0% |
| Mental health outpatient facilities (per 100,000) | 88.0% | 88.0% | 88.0% | 91.3% |
| Mental health units in general hospitals (per 100,000) | 84.0% | 88.0% | 88.0% | 91.3% |
| Mental health units in general hospitals admissions (per 100,000) | 96.0% | 96.0% | 96.0% | 100.0% |
| Mental hospital admissions (per 100,000) | 96.0% | 96.0% | 96.0% | 95.7% |
| Nurses in mental health sector (per 100,000) | 84.0% | 88.0% | 88.0% | 91.3% |
| Outpatient visits (per 100,000) | 96.0% | 96.0% | 96.0% | 100.0% |
| Psychiatrists in mental health sector (per 100,000) | 84.0% | 88.0% | 88.0% | 91.3% |
| Psychologists in mental health sector (per 100,000) | 84.0% | 88.0% | 88.0% | 91.3% |
| Schizophrenia (DALYs) | 12.0% | 12.0% | 12.0% | 13.0% |
| Schizophrenia (Prevalence) | 96.0% | 96.0% | 96.0% | 95.7% |
| Social workers in mental health sector (per 100,000) | 84.0% | 88.0% | 88.0% | 91.3% |
| Self-reported life satisfaction | 56.0% | 56.0% | 56.0% | 52.2% |
| Share of people who say they are happy | 84.0% | 84.0% | 84.0% | 87.0% |

DALYs: Disability-Adjusted Life Years.

Even for core indicators such as disorder prevalence and DALYs, data availability was uneven across income groups, while subjective well-being indicators showed moderate but still substantial levels of missingness. These patterns highlight structural limitations in global mental health surveillance and directly constrain the interpretability of longitudinal ecological models applied in this context.

## 4. Discussion

The aim of science is human progress; however, in a vast sea of data and publications that is in exponential expansion, it has become difficult to answer the question of how useful global research really is, in this case, on mental health [9,20,21]. It is expected that when more research is produced on a topic, the population-level indicators related to it will also improve [2]. However, this question, to our knowledge, has not been properly addressed, since no formal methodology has been developed with that aim. It is theoretically possible to estimate this coherence in science, given that we usually have access, over the years, to both the scientific production and the population-level indicator data [9]. This study examined the behaviour of longitudinal ecological models linking scientific production to population-level indicators, using global mental health research as an empirical case. Interpreted through a methodological lens, the findings provide insight not into the effectiveness of mental health research per se, but into the conditions under which such models generate interpretable associations, and the circumstances in which they fail to do so.

One of the most salient observations emerging from the analyses is the marked divergence in association patterns across income groups. In higher-income settings, inverse associations between publication volume and several mental disorder indicators were consistently observed, whereas coefficients in low-income countries tended to follow the opposite direction. From a methodological standpoint, this contrast should not be interpreted as evidence of beneficial or harmful effects of research activity. Rather, it reflects how ecological models respond to structurally distinct contexts characterized by differences in research capacity, health system maturity, and data generation processes [22].

These findings underscore that income stratification is not merely a descriptive convenience, but a critical analytical dimension that fundamentally shapes model outputs [22]. Aggregating countries across income levels would have masked this heterogeneity and potentially produced misleading pooled estimates. The observed divergence therefore reinforces the necessity of stratified analyses when applying longitudinal ecological frameworks to global datasets [23].

The extreme heterogeneity identified in the meta-analytic stage, reflected by I^2^ values approaching or exceeding 90% for most indicators, represents a central methodological result. Rather than indicating analytical failure, this degree of heterogeneity highlights the limits of summarizing association patterns across contexts that differ structurally in data availability, indicator validity, and surveillance intensity [24].

The fact that only one subjective well-being indicator retained a statistically significant pooled estimate further illustrates this point. Pooled effects in such settings should be interpreted as descriptive summaries of highly variable associations, not as stable parameters. The inability of meta-regression to meaningfully explain between-group heterogeneity reinforces the conclusion that much of the observed variability arises from unmeasured structural factors rather than from modifiable covariates included in the models [25].

The extensive missingness observed across mental health indicators constitutes another key methodological finding. Infrastructure-related indicators and government expenditure measures were unavailable for large portions of the study period, particularly in lower-income strata [19]. Importantly, this missingness is not random; it reflects longstanding limitations in global mental health surveillance systems.

Within longitudinal ecological models, such patterns directly constrain inferential capacity. Associations cannot be estimated where indicators are sparsely measured, and where they are estimated, their interpretability is contingent on the stability and completeness of the underlying data. Treating missingness as an informative feature rather than a technical nuisance allowed these limitations to be surfaced explicitly, rather than obscured through imputation or aggregation strategies that could artificially inflate model performance [19].

Taken together, these findings suggest that longitudinal ecological models linking scientific production to population-level indicators are best understood as exploratory tools rather than evaluative instruments. They are capable of identifying broad association patterns and highlighting structural asymmetries, but they are poorly suited for attributing changes in population outcomes to research activity.

From a meta-research perspective, the results caution against simplistic interpretations of research impact based on aggregate associations. Apparent coherence, or incoherence, between scientific output and population indicators (scientific coherence) may reflect data infrastructure, reporting practices, or contextual heterogeneity as much as any underlying relationship between knowledge production and societal outcomes [9,26,27]. Consequently, assessments of research coherence should incorporate explicit consideration of indicator coverage, contextual stratification, and model sensitivity [5].

### 4.1. Strengths and methodological limitations

A key strength of this study lies in its integration of large-scale bibliometric data with multiple domains of population-level indicators over an extended temporal horizon. The sequential analytical framework enabled systematic examination of association patterns, heterogeneity, and potential moderators while maintaining transparency at each stage.

Nonetheless, some limitations must be acknowledged. The ecological nature of the design precludes causal inference, and publication volume represents an imperfect proxy for scientific activity, capturing quantity but not quality, relevance, or translational uptake. Moreover, the uneven availability of mental health indicators limits both the scope and interpretability of the analyses, particularly in low-resource settings.

These limitations are not unique to this study; rather, they reflect broader structural constraints inherent to global research evaluation efforts. Explicitly articulating them is therefore essential for contextualizing findings and guiding future methodological developments.

### 4.2. Future methodological directions

Future work in this area may benefit from combining ecological approaches with complementary methodologies, such as case-based analyses, quasi-experimental designs, or linkage studies integrating policy implementation data. Expanding and standardizing global mental health surveillance systems would substantially enhance the interpretability of longitudinal models and enable more nuanced assessments of research-outcome alignment [28].

Importantly, methodological innovation should proceed alongside conceptual clarity regarding what such models are intended to measure, and what they are not. Without this clarity, there is a risk of overinterpreting descriptive associations as evidence of effectiveness or failure.

## 5. Conclusions

This study provides an empirical assessment of the methodological performance of longitudinal ecological models that link scientific production to population-level indicators, using global mental health research as a case study. When examined through a meta-research lens, the findings demonstrate that such models are capable of identifying broad association patterns and structural asymmetries across contexts, but that their interpretability is strongly constrained by heterogeneity, data availability, and contextual stratification.

The pronounced divergence in associations across income groups, the persistence of extreme between-group heterogeneity, and the limited explanatory capacity of meta-regression collectively indicate that aggregate links between research output and population-level indicators cannot be interpreted as proxies for research effectiveness or societal impact. Rather, these patterns reflect how ecological models respond to uneven research capacity, surveillance infrastructure, and indicator coverage at the global level.

Importantly, the results underscore that missing data and inconsistent indicator measurement are not peripheral technical issues but central methodological determinants of model behaviour. In settings where population-level indicators are sparse or irregularly collected, longitudinal ecological approaches are inherently limited in their ability to generate stable or generalizable associations, regardless of analytical sophistication.

From a methodological standpoint, these findings suggest that longitudinal ecological models should be used with caution in research evaluation contexts. Their primary value lies in exploratory analysis, hypothesis generation, and the identification of structural gaps in data systems, rather than in the attribution of changes in population outcomes to scientific activity. Explicit recognition of these limits is essential to avoid overinterpretation and to ensure responsible use of such models in meta-research and science policy discussions.

Future methodological advances in this area will depend not only on improved analytical techniques but also on strengthened global surveillance systems and clearer conceptual frameworks for assessing research coherence. Aligning scientific production with population-level indicators requires both robust data infrastructures and methodological approaches that are explicitly calibrated to the questions they are intended to address.

## Abbreviations

DALYs: Disability-Adjusted Life Years

## Declarations

### Author contributions

AAAM, DAHP, FJVL, and IDLM contributed to the conception and design of the study. AAAM, DAHP, FJVL, IDLM and SK carried out data collection and data cleaning. AAAM, DAHP, and IDLM performed the statistical analyses and were involved in data organization and presentation. AAAM, DAHP, FJVL, IDLM and SK drafted the manuscript. AAAM, DAHP, FJVL, IDLM and SK critically reviewed the manuscript with suggestions for improvement and revision. All authors read and approved the final version.

## Acknowledgments

None.

## Ethics approval and consent to participate

Not applicable.

## Consent for publication

Not applicable.

## Clinical trial number

Not applicable.

## Availability of data and materials

The dataset generated and analyzed is available at https://doi.org/10.5281/zenodo.18417489.

## References

1. Ninkov A, Frank JR, Maggio LA. Bibliometrics: Methods for studying academic publishing. Perspect Med Educ. 2022 Jun;11(3):173–176. doi: 10.1007/s40037-021-00695-4

2. Lozada-Martinez ID, Hernandez-Paz DA, Fiorillo-Moreno O, Picón-Jaimes YA, Bermúdez V. Meta-research in biomedical investigation: gaps and opportunities based on meta-research publications and global indicators in health, science, and human development. Publications. 2025; 13(1):7. 10.3390/publications13010007

3. Wolfenden L, Mooney K, Gonzalez S, Hall A, Hodder R, Nathan N, et al. Increased use of knowledge translation strategies is associated with greater research impact on public health policy and practice: an analysis of trials of nutrition, physical activity, sexual health, tobacco, alcohol and substance use interventions. Health Res Policy Syst. 2022 Jan 31;20(1):15. doi: 10.1186/s12961-022-00817-2.

4. Greenhalgh T, Raftery J, Hanney S, Glover M. Research impact: a narrative review. BMC Med. 2016 May 23;14:78. doi: 10.1186/s12916-016-0620-8

5. Chan SL, Ho CZH, Khaing NEE, Ho E, Pong C, Guan JS, et al. Frameworks for measuring population health: A scoping review. PLoS One. 2024 Feb 13;19(2):e0278434. doi: 10.1371/journal.pone.0278434

6. Sarquis Rivera MA, Hernandez-Paez DA, Galván-Barrios J, Barceló-Martinez E, Narvaez-Rojas AR, Lozada-Martinez ID. Exploring the potential impact of medical errors research on population health. PLoS One. 2026 Mar 12;21(3):e0340153. doi: 10.1371/journal.pone.0340153

7. Revolledo Caicedo MA, Montoya Obando AF, Hernández-Páez DA, Fiorillo-Moreno O, Galván-Barrios J, Delgado P, et al. Research on medical errors exhibits diverse associations with global indicators of science, development, and health across geographic regions: A scientometrics study. Medicine (Baltimore). 2025 Jul 18;104(29):e42985. doi: 10.1097/MD.0000000000042985

8. Romero-Freile E, Ortiz-Cerchar D, Hernández-Páez DA, Fiorillo-Moreno O, Picón-Jaimes YA, Beltrán-Venegas T, et al. Six Sigma research in healthcare is associated with global health, research and development indicators: A scientometrics study based on countries by income groups. An Sist Sanit Navar. 2025 Aug 28;48(2):e1124. doi: 10.23938/ASSN.1124

9. Hernandez-Paez DA, Acuña-Rodríguez M, Pertuz-López NM, Visconti-Lopez FJ, Martinez-Royert JC, Lozada-Martinez ID. Limited scientific coherence between global mental health research and indicators of science, health, mental health, and society: a longitudinal analysis across world regions. Front Psychol. 2026; 16: 1649735. doi: 10.3389/fpsyg.2025.1649735

10. Purtle J, Nelson KL, Counts NZ, Yudell M. Population-Based Approaches to Mental Health: History, Strategies, and Evidence. Annu Rev Public Health. 2020 Apr 2;41:201–221. doi: 10.1146/annurev-publhealth-040119-094247

11. Ioannidis JP, Fanelli D, Dunne DD, Goodman SN. Meta-research: Evaluation and Improvement of Research Methods and Practices. PLoS Biol. 2015 Oct 2;13(10):e1002264. doi: 10.1371/journal.pbio.1002264

12. Burg D, Ausubel JH. Trajectories of COVID-19: A longitudinal analysis of many nations and subnational regions. PLoS One. 2023 Jun 23;18(6):e0281224. doi: 10.1371/journal.pone.0281224

13. The World Bank. World Bank Country and Lending Groups. Jan 25, 2026. https://datahelpdesk.worldbank.org/knowledgebase/articles/906519-world-bank-country-and-lending-groups

14. World Health Organization. The Global Health Observatory. Jan 25, 2026. https://www.who.int/data/gho

15. The World Bank. Indicators. Jan 25, 2026. https://data.worldbank.org/indicator

16. Our World in Data. Data. Jan 25, 2026. https://ourworldindata.org/search

17. Institute for Health Metrics and Evaluation. Research and analysis. Jan 25, 2026. https://www.healthdata.org/research-analysis

18. Zhao Y, Ding-Geng C. Statistical Modeling in Biomedical Research: Contemporary Topics and Voices in the Field. 1st edition. Springer Cham. Switzerland. doi: 10.1007/978-3-030-33416-1

19. Heymans MW, Twisk JWR. Handling missing data in clinical research. J Clin Epidemiol. 2022 Nov;151:185–188. doi: 10.1016/j.jclinepi.2022.08.016

20. Cruz Rivera S, Kyte DG, Aiyegbusi OL, Keeley TJ, Calvert MJ. Assessing the impact of healthcare research: A systematic review of methodological frameworks. PLoS Med. 2017 Aug 9;14(8):e1002370. doi: 10.1371/journal.pmed.1002370

21. Thonon F, Boulkedid R, Delory T, Rousseau S, Saghatchian M, van Harten W, et al. Measuring the outcome of biomedical research: a systematic literature review. PLoS One. 2015 Apr 2;10(4):e0122239. doi: 10.1371/journal.pone.0122239

22. Aggarwal M, Hutchison B, Wong ST, Katz A, Slade S, Snelgrove D. What factors are associated with the research productivity of primary care researchers in Canada? A qualitative study. BMC Health Serv Res. 2024 Mar 1;24(1):263. doi: 10.1186/s12913-024-10644-6

23. Shimonovich M, Campbell M, Thomson RM, Broadbent P, Wells V, Kopasker D, et al. Causal Assessment of Income Inequality on Self-Rated Health and All-Cause Mortality: A Systematic Review and Meta-Analysis. Milbank Q. 2024 Mar;102(1):141–182. doi: 10.1111/1468-0009.12689

24. Heyard R, Held L. Meta-regression to explain shrinkage and heterogeneity in large-scale replication projects. PLoS One. 2025 Aug 1;20(8):e0327799. doi: 10.1371/journal.pone.0327799

25. Jansen JP, Cope S. Meta-regression models to address heterogeneity and inconsistency in network meta-analysis of survival outcomes. BMC Med Res Methodol. 2012 Oct 8;12:152. doi: 10.1186/1471-2288-12-152

26. Arsalan MH, Mubin O, Mahmud AA, Perveen S. What factors influence research impact? An empirical study on the interplay of research, publications, researchers, institutions, and national conditions. 2025; 10(1):188–227. doi: 10.2478/jdis-2025-0001

27. Cao Z, Zhang L, Wang Z, Li C, Sivertsen G. How does scientific research generate impact beyond academia? Cross-disciplinary comparison based on REF impact cases. Humanit Soc Sci Commun. 2025; 12: 1856. doi: 10.1057/s41599-025-06129-4

28. Benjet C. Improving mental health with more equitable surveillance data. Lancet Glob Health. 2025 May;13(5):e785–e786. doi: 10.1016/S2214-109X(25)00052-X

